# Can mindfulness meditation alter consciousness? An integrative interpretation

**DOI:** 10.1101/024174

**Authors:** Jordi Manuello, Ugo Vercelli, Andrea Nani, Tommaso Costa, Franco Cauda

**Author notes:** **Corresponding author:** Dr. Andrea Nani.

## Abstract

Mindfulness meditation has been practiced in the East for more than two millennia, but in last years also Western neurscientists drown their attention to it. Mindfulness basically refers to moment to moment awareness. In this review we summarize different hypotheses concerning effects of mindfulness meditation practice and cerebral correlates accounting for these; furthermore we expose some of the most relevant theories dealing with different aspects of consciousness. Finally we propose an integration of mindfulness meditation with consciousness, supported by the identification of brain areas involved in both of them, namely Anterior Cingular Cortex (ACC), Posterior Cingular Cortex (PCC), Insula and Thalamus.

## 1. INTRODUCTION

Meditation is a practice that has existed for many centuries. It involves different techniques and can be found in a variety of cultural traditions, ranging from Indian and Chinese to the Arab and Western worlds. However, meditation has traditionally been associated with the Eastern culture and spirituality, especially with the Indian religion of Hinduism – whose ancient scriptures (The Vedas) report the earliest references to this practice – and the philosophy of Buddhism, which holds meditation as a key part of its doctrine (Siegel et al., 2008).

In recent years Western societies have become more accustomed to meditation, in particular through the interest that Buddhism has attracted by virtue of the charismatic figure of the current Dalai Lama Tenzin Gyatso. Moreover, meditative practices have been investigated by a number of scientific studies, whose findings have gained increasing attention in healthcare treatment programs within psychotherapeutic contexts (Samuel, 2014; Tang et al., 2015).

Although meditation escapes a univocal definition, it is nonetheless possible to intuitively deduce what it is by saying what it is not. Meditation is neither a method for clearing the mind nor a method for reaching emotional imperturbability. It is not a way to pursue a state of beatitude or a way to avoid sorrow and pain (Siegel et al., 2008). Nor does it imply a secluded life.

Often the meditative state is improperly associated with esotericism and mysticism. But as the Theravada monk Nyanaponika Thera (1998) clearly highlights: “Mindfulness […] is not at all a ‘mystical’ state, beyond the ken and reach of the average person. It is, on the contrary, something quite simple and common, and very familiar to us. In its elementary manifestation, known under the term ‘attention’, it is one of the cardinal functions of consciousness without which there cannot be perception of any object at all.” (Thera, 1962). As we shall see, this position places meditation directly in the spotlight of neuroscience.

Although there are many different meditation techniques, all of them share the fundamental aspect of “sati”, a Pali word translated into English as “mindfulness” for the first time in 1921 (Awasthi, 2012; Siegel et al., 2008). Sati is also a core concept of Buddhist philosophy. Jon Kabat-Zinn, who pioneered the mindfulness approach within the therapeutic context, defines this state of mind as “the awareness that emerges through paying attention on purpose, in the present moment, and nonjudgmentally to the unfolding of experience moment to moment”(Kabat-Zinn, 2003).

This review aims to integrate the findings of studies that have used functional Magnetic Resonance Imaging (fMRI) to investigate the morphological and functional modifications observed in people practicing meditation with what neuroscientists have thus far discovered about the neural processes that promote the emergence and maintenance of consciousness.

## 2. DIFFERENT STYLES OF MEDITATION

According to Siegel (2008) we can distinguish three meditative techniques within the general framework of “Mindfulness Based Meditation” (MBM).

### Concentration meditation

This technique is based on focusing on a specific object, such as the breath or a mantra. The guideline is to gently bring the mind back to the focal object whenever you notice that you are wandering. The Pali term for this technique is “Samatha bhavana”, which can be translated into English as “to foster concentration”.

### Mindfulness meditation

This technique does not use a focal object but rather encourages the exploration of the ever-changing experience as it unfolds through time. The guideline is to pay attention to whatever flickers across consciousness from one moment to the next. The Pali term for this technique is “Vipassana bhavana”, which translates as “to foster interior awareness”.

### Loving-kindness meditation

With this technique the mind is led to concentrate on gentle statements such as “May I and all the other creatures be safe, happy, healthy and live in simplicity”. The aim is to soften emotions and observe the experience with a non-judgmental attitude, free from overwhelming emotionality. The Pali term for this technique is “Metta bhavana”, which translates as “to foster fondness”.

Even though they are distinct, these three techniques can be used together; in fact they all foster “sati” and, at the same time, require it to be continuously reinforced in a sort of circular mental process.

## 3. MEDITATION AND THE BRAIN

Since its early stages, meditation has been thought to be the primary method for both enhancing awareness and keeping the body and mind in good health (Siegel et al., 2008). It is therefore not surprising that over the last few years Mindfulness-Based Interventions (MBIs), which are therapeutic approaches based on MBM, have attracted increasingly more interest in a variety of fields, ranging from psychology and neuroscience to public health and education circles (Chiesa and Serretti, 2010; Hölzel et al., 2011). Mindfulness-Based Stress Reduction (MBSR), Mindfulness-Based Cognitive Therapy (MBCT) and Integrative Body-Mind Training (IBMT) are the most renowned MBI techniques. In particular MBSR, which was devised in 1979 at the University of Massachusetts Medical Center (Kabat-Zinn, 2003), is currently used as an alternative or integrative clinical approach for the treatment of psychological issues in people with chronic diseases (Chiesa and Serretti, 2011; Merkes, 2010). Still, the neuroanatomical and functional correlates that underpin the beneficial effects of MBIs are yet to be fully understood (Tang et al., 2015).

In spite of the different styles of meditation and the assortment of MBIs, the aspect of “sati” or “mindfulness” is the common thread that unites them all. As we have seen, a state of mindfulness is characterized by consciously paying attention to the unfolding experience of the present moment (Kabat-Zinn, 2003). Accordingly, since mindfulness directly involves both consciousness and attention, the neural correlates of these brain processes and those of meditative states should manifest strong similarities.

Interoceptive Attention (IA) has been put forward as a crucial process in order to account for mindfulness meditation. Interoception is composed of several bodily sensations related to digestion, blood circulation, breathing and proprioception (Farb et al., 2013).

Neuroanatomical studies have provided evidence of spino-thalamo-cortical pathways projecting to the granular posterior insula and medial agranular insula areas, which are thought to function as the primary interoceptive cortex (Flynn, 1999). Moreover, descending projections toward sensory and motor areas of the brainstem originate from the insula and the anterior cingulate cortex (ACC) (Craig, 2009a).

In line with these neuroanatomical findings, a recent experiment by Farb et al., (2013) found that, after 8 weeks of MBSR, participants were showing an increased functional plasticity in the medial and anterior insula, areas which are associated with the awareness of the present moment (Craig, 2009a; Farb et al., 2007). What is more, the practice of mindfulness meditation might promote functional connectivity between the posterior insula and the anterior insular gyrus, thus increasing the overall activation of the anterior insula and, at the same time, reducing the recruitment of the dorsomedial prefrontal cortex (DMPFC) (Farb et al., 2013). DMPFC deactivation has also been found in conjunction with exogenous stimulation of interoceptive signaling pathways, e.g. during gastric distension (Van Oudenhove et al., 2009). On the contrary, DMPFC activation is associated with the executive control of behavior related to the shift of attention during performances of problem solving performances (Mullette-Gillman and Huettel, 2009) and, possibly, with either stimulus-independent or stimulus-oriented thought in mind-wandering (Christoff et al., 2009). Therefore, DMPFC deactivation after MBSR training might be one of the signs that can help to distinguish between mindfulness and mind-wandering as well as between mindfulness and mental effort (Farb et al., 2010; Farb et al., 2007).

To evaluate the impact of mindfulness meditation practice, a recent study compared MBSR with an aerobic exercise for stress reduction. The results showed that only MBSR significantly helps to regulate negative emotions in people with social anxiety. According to the authors, such effect might be due to the functional integration of different brain networks, which occurs during somatic, attentional and cognitive control (Goldin et al., 2013).

Other investigations have focused on whether the practice of long-term meditation could cause brain structural changes, suggesting that long-term meditation might be associated with increased cortical thickness, especially in the prefrontal and right anterior insular cortices, which are involved in attention, interoception, and sensory data elaboration (Lazar et al., 2005; Sato et al., 2012). Significantly, one study was able to classify participants as meditators or non-meditators on the basis of subtly different patterns across the brain (Sato et al., 2012). Using a multivariate pattern recognition method, such as the Support Vector Machine (SVM), this study examined whether or not a single subject could be identified as a regular meditator. The SVM had an accuracy of 94.87%, allowing the identification of 37 out of 39 participants. The right precentral gyrus, the left entorhinal cortex, the right pars opercularis cortex, the right basal putamen, and the thalamus bilaterally were the most informative brain regions used in the classification. The involvement of these areas strongly suggests the potential of mindfulness meditation to increase awareness and recognition of bodily sensations, as well as to improve interoceptive observational skills (Kozasa et al., 2012; Lazar et al., 2005).

## 4. THE NEUROSCIENCE OF CONSCIOUSNESS

As we have seen, the concepts of mindfulness and consciousness are inextricably intertwined. Both neurophysiological and neuroimaging studies have provided evidence that the neural correlates of consciousness can be described in virtue of a bidimensional model, based on the parameter of the level of arousal on the one hand, and on the parameter of the intensity of the different contents of experience on the other (Cavanna et al., 2011; Laureys, 2005; Laureys et al., 2004; Nani et al., 2013).Within this framework, the arousal dimension evaluates the quantitative features of consciousness, whereas the content dimension addresses the qualitative features of subjective awareness (Blumenfeld, 2009; Plum and Posner, 1980; Zeman, 2001). In other words, the level of arousal specifies the degree of wakefulness, which can range from full alertness through drowsiness and sleep to coma (Baars et al., 2003; Laureys and Boly, 2008). The integrity of the ascending ponto-mesodiencephalic reticular pathways and widespread thalamo-cortical networks is essential for promoting and maintaining consciousness (Steriade, 1996a, b).

The experiential contents are all the things that can pop up in the conscious mind, such as sensations, emotions, thoughts, memories, intentions, etc. They are likely to be caused by the interaction between exogenous factors (i.e., environmental stimuli) and endogenous factors (i.e., internal bodily inputs). The dimension of contents can therefore be divided into external awareness (what we perceive through the senses) and internal awareness (thoughts which are independent of specific external stimuli) (Demertzi et al., 2013) [**Fig. 1**].

**Figure 1.**
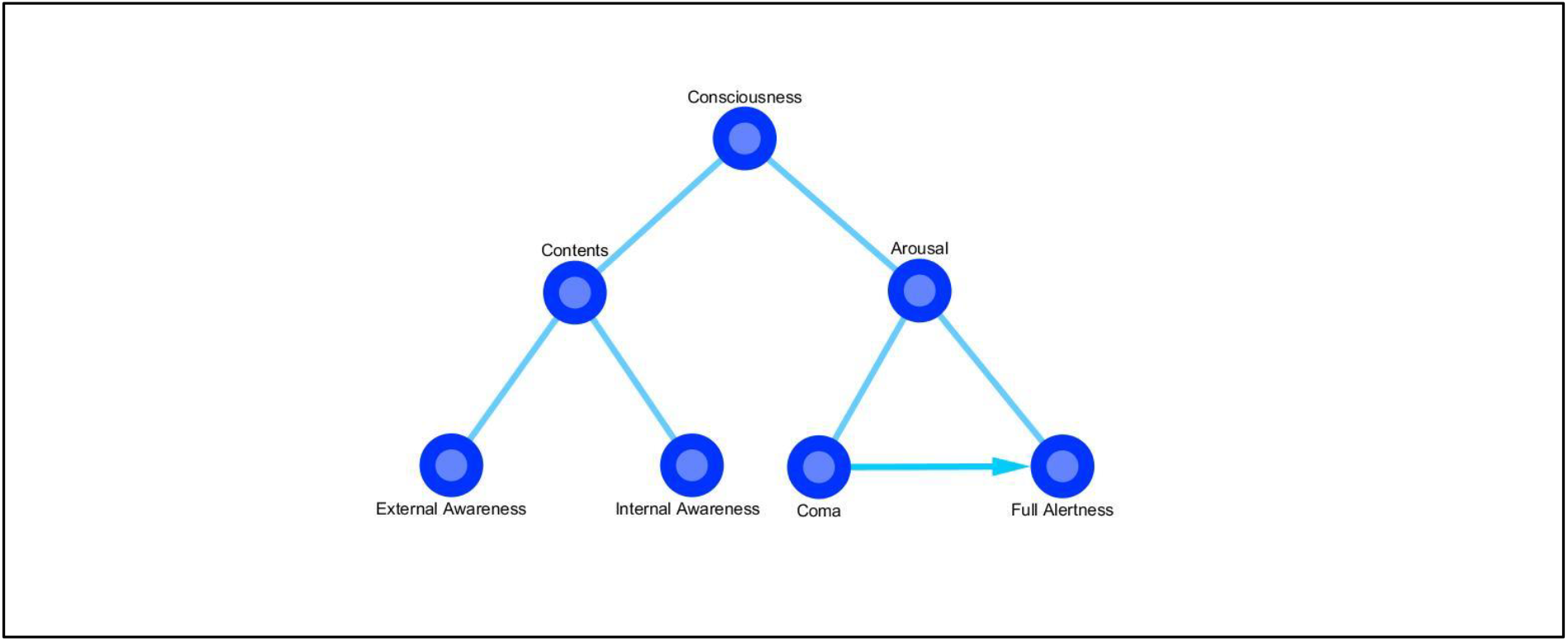
A bidimensional model of consciousness. According to a bidimensional model, the neural correlates of consciousness can refer to both the level of arousal (ranging from full alertness to coma) and the different contents of experience, which can in turn be divided into external awareness and internal awareness.

This distinction is important because internal and external awareness seem to involve different neural correlates. Demertzi et al. (2013) described an “internal awareness network”, which includes the posterior cingulate cortex (PCC), the ACC, the precuneus and the medial prefrontal cortex (MPFC), and an “external awareness network”, which includes the dorsolateral prefrontal cortex (DLPFC) and the posterior parietal cortex (PPC).

The interaction of these two networks creates what has been called a “global neuronal workspace”, which is thought to be of fundamental importance in supporting consciousness (Baars et al., 2003; Dehaene and Changeux, 2011). What is more, internal and external awareness networks have been shown to overlap with some of the areas involved in the Default Mode Network (DMN), such as the PCC, precuneus and MPFC, as well as with some of the areas involved in the Salience Network (SN), such as the ACC and the thalamus, and in the Central Executive Network (CEN), such as the DLPFC and the PPC.

### 4.1. Consciousness and self-consciousness

Within the neuroscientific study of consciousness, other important and debated issues are the origin of self, the construction of self-awareness and the relationship between consciousness and self-consciousness. The concept of self is as difficult to define as the concept of consciousness. Most research (Metzinger and Gallese, 2003; Pacherie, 2008; Roessler and Eilan, 2003), focusing on the central representation of different parts of the body, has linked the sense of self to other concepts, such as *agency* – i.e., “the sense that a person’s action is the consequence of his or her intention” (Seth et al., 2012) – and *embodiment* – i.e., “the sense of being localized within one’s physical body” (Arzy et al., 2006). Agency and embodiment might be connected with what has been called the “minimal phenomenal selfhood” (MPS), that is, “the experience of being a distinct, holistic entity capable of global self-control and attention, possessing a body and a location in space and time” (Blanke and Metzinger, 2009). The MPS could be disrupted in brain injured patients, who are likely to be subject to autoscopic experiences (Blanke et al., 2004; Blanke and Mohr, 2005; Brugger, 2006; Devinsky et al., 1989).

A framework based on the concept of *agency* in association with interoceptive predictive coding has been put forward in order to account for the feeling of *conscious presence*, which has been defined as “the subjective sense of reality of the world and of the self within the world” (Seth et al., 2012). This model is characterized by predictive signals of agency and relies on a mechanism of interoceptive prediction error implemented in the perception of the state of the body through autonomic physiological responses, which are commonly involved in the generation of emotions (Craig, 2009b; Critchley et al., 2004). The mechanism of interoception was traditionally considered to be bound to visceral sensations only, but contemporary neuroanatomical and neurophysiological research suggests that it may also involve information coming from the muscles, articulations, skin, and organs. And all this various information appears to be conjointly processed (Cauda et al., 2012; Craig, 2002).

According to this model, the sense of conscious presence emerges when interoceptive prediction signals and real input signals match, so that the error signals are suppressed (Seth et al., 2012) [**Fig. 2**].

**Figure 2.**
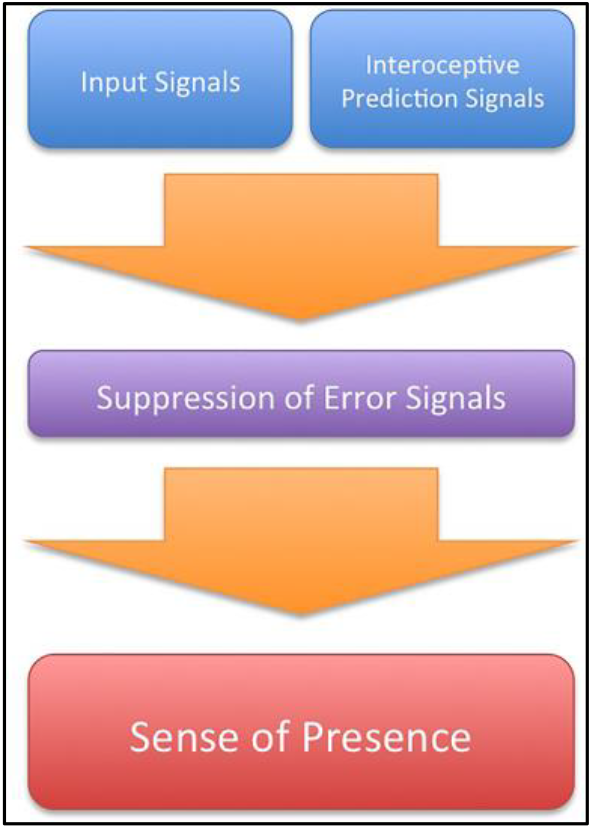
Schematic model of the sense of presence. When interoceptive prediction signals and input signals match, the error signals are suppressed and the sense of presence emerges (adapted from Seth et al., 2012).

The cortical areas thought to be fundamentally involved in this process are the orbitofrontal cortex, the ACC and the insula (Critchley et al., 2004); it has been proposed that the insula, in particular, operates the integration between interoceptive and exteroceptive signals, thus promoting the generation of subjective emotional states (Cauda et al., 2011; Seth et al., 2012).

Intriguingly, the anterior insula and the ACC are among the few human brain areas in which von Economo neurons (VENs) are present (Craig, 2004; Sturm et al., 2006; von Economo, 1926, 1927; von Economo and Koskinas, 1925). These large spindle-shaped neurons have been supposed to be involved in the perception of bodily states (Allman et al., 2005; Cauda et al., 2014). What is more, they have recently been associated with the neural correlates of consciousness on the basis of two principal morphological and cytochemical findings (Cauda et al., 2014; Cauda et al., 2013; Critchley and Seth, 2012; Medford and Critchley, 2010; Menon and Uddin, 2010). First, consciousness is likely to be supported by long-range connection (Cauda et al., 2014; Dehaene and Changeux, 2011; Dehaene et al., 1998) and VENs are indeed long projection neurons. Second, VENs selectively express high levels of bombesin-related peptide neuromedin B (NMB) and of gastrin releasing peptide (GRP), which are “involved in the peripheral control of digestion and also known to participate in the conscious awareness of bodily states” (Allman et al., 2010, 2011; Cauda et al., 2014; Stimpson et al., 2011).

Within the framework of Seth’s model, VENs might project to the visceral autonomic nuclei (i.e., the periaqueductal gray and the parabrachial nucleus), which are significantly involved in interoception (Allman et al., 2005; Butti et al., 2009; Cauda et al., 2014; Craig, 2002; Seeley, 2008). The anterior insula and the ACC, which are functionally (Taylor et al., 2009; Torta and Cauda, 2011) and structurally (van den Heuvel et al., 2009) interconnected, are part of the SN (Medford and Critchley, 2010; Palaniyappan and Liddle, 2012; Seeley et al., 2007b). This network responds to behaviorally salient events and things, by identifying the relevant aspects or qualities by which they stand out relative to the surrounding environment. Therefore, it seems plausible that the SN might play a crucial role in Seth’s model, by processing exteroceptive signals with a certain degree of salience (Seth et al., 2012). Moreover, recent evidence suggests that a specific part of the SN (i.e., the anterior insula) could induce a switch between the CEN and the DMN, thus orienting attention towards the external or internal world (Bressler and Menon, 2010).

### 4.2. Consciousness and the predicting brain

Another hypothesis in which consciousness of the present moment relies heavily on neurofunctional mechanisms for making predictions was put forward by Moshe Bar (2007). His “proactive brain” theory holds that the brain continuously makes predictions on the basis of sensory and cognitive information. Bar’s hypothesis is supported by observing that most of the DMN, which is active during the resting state (Tang et al., 2012), overlaps with the brain areas (the MPFC, medial parietal cortex and medial temporal lobe) that are recruited during the performance of tasks requiring associative elaboration (Bar et al., 2007).

A similar view of the brain architecture can be found in the “Bayesian brain” hypothesis, according to which “we are [always] trying to infer the causes of our sensations based on a generative model of the world.” (Dayan et al., 1995; Friston, 2012; Gregory, 1980; Kersten et al., 2004; Knill and Pouget, 2004; Lee and Mumford, 2003). As a consequence, we commonly try to predict the future by taking into account the statistical history of past events and stimuli (Bar, 2007).

All these predictive theories (Seth’s model, the “proactive brain” and “Bayesian brain” hypotheses) could be reappraised within the more general context of the “free energy principle” (Friston et al., 2006), according to which “any self-organizing system that is at equilibrium with its environment must minimize its free energy” (Friston, 2010). Free energy can be thought of as a measure of the difference between the distribution of environmental energy that acts on biological systems and the distribution of energy that is embodied in the organization of those biological systems. In other words, free energy emerges from the exchanges of energy between the biological systems and their environment (Friston et al., 2006). Therefore, if individuals are to be thought of as the sum of their models of the world, they need to find a point of equilibrium in which free energy is minimized. And the emergence of consciousness seems to be a very suitable way to reach and maintain this equilibrium.

### 4.3 The global workspace theory of consciousness

As we have seen in the previous paragraphs, the large spindle-shaped VENs could play an important role not only in predictive models of brain functioning but also in theories that aim to account for the emergence of consciousness. In particular, VENs could be pivotal in the “Global Workspace Model” of conscious elaboration (Baars, 1988; Dehaene and Changeux, 2011). This model assumes two different computational spaces within the brain (Dehaene et al., 1998). One space is a network composed of several functionally specialized modular subsystems (Baars, 1988; Shallice, 1988). Each subsystem resides in a specific cortical region and has medium-range connections to other areas (Mesulam, 1998). The second space is a distributed global workspace (GW) composed of cortical neurons, reciprocally connected through horizontal long-range bidirectional projections, and whose concentration is variously related to different brain regions. These long-range projections could easily explain the property of reportability (Weiskrantz, 1997), which is a characteristic feature of conscious phenomena. In fact, within the GW both speech and motor areas can be connected to the associative areas that deal with the contents of experience.

According to the model, “what we subjectively experience as a conscious state” is the distributed availability of information within the common global space, which is made possible by the long-range neuronal projections (Dehaene and Naccache, 2001). As a consequence, conscious stimuli seem to be less encapsulated in specific processes than unconscious ones (Dehaene and Changeux, 2011). Moreover, evidence shows that the GW becomes active during non-routinized tasks, it gradually switches off during the process of learning, while it suddenly activates again in case of error detection (Dehaene et al., 1998). From the neuroanatomical perspective, the brain areas that are likely to be associated with the GW are the dorsolateral prefrontal cortex and the ACC (Dehaene et al., 1998), which are therefore thought to be involved in the conscious awareness of subjective states (Grafton et al., 1995; Sahraie et al., 1997).

## DISCUSSION

The practice of mindfulness meditation can have effects in terms of increased attention, control and orientation, along with improvements in cognitive flexibility. Many practitioners describe what they feel during meditation as “undistracted awareness” and “effortless doing” (Garrison et al., 2013). Accordingly, Tang et al. (2012) observed that the effort needed to maintain attention tends to fade during a meditative session.

If the hypothesis that mindfulness meditation can have an impact on consciousness is correct, we would expect some degree of overlap to exist between the brain areas involved in both processes and, consequently, a change in the activity of those areas in (at least) long-term meditators. In line with this hypothesis, contemporary research has highlighted that some main brain areas are strictly associated with both meditation and consciousness [**Fig. 3**] [**Fig. 4**].

**Figure 3.**
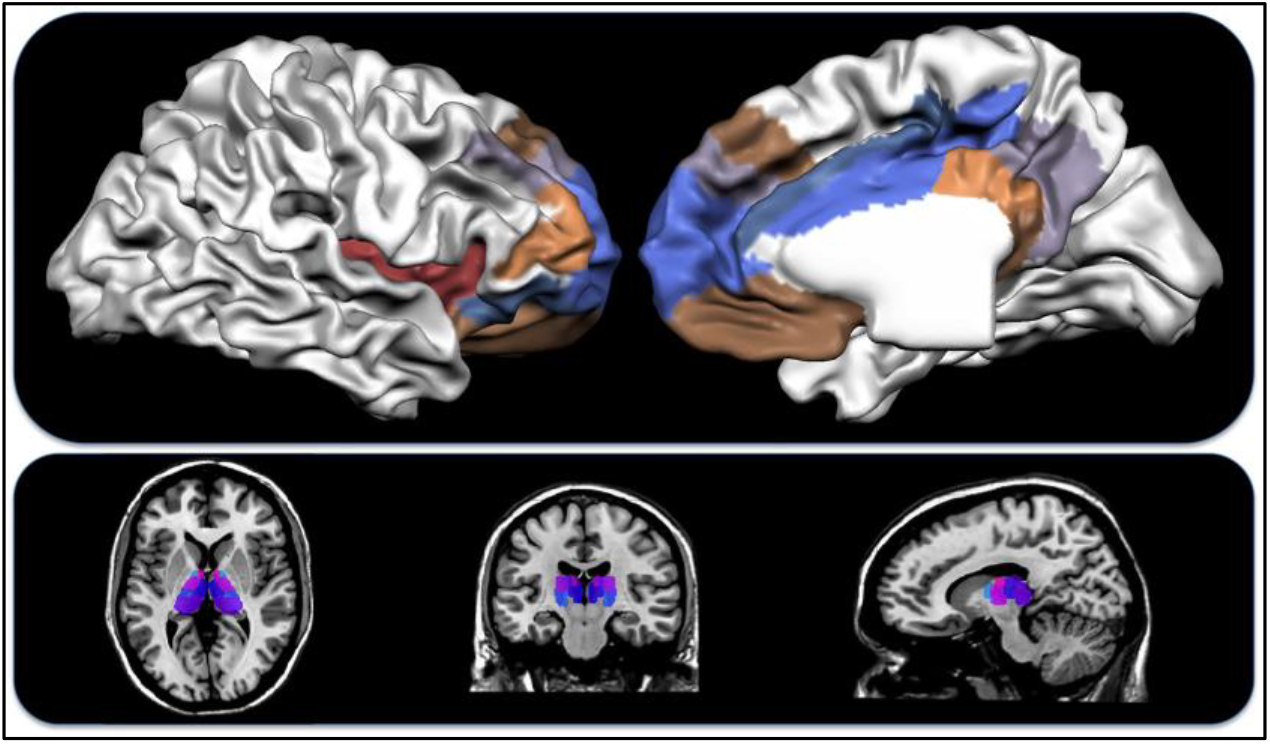
Brain areas involved in both mindfulness meditation and consciousness. Top: the insular cortex and prefrontal lateral areas (left), medial areas (right). Bottom: the thalamus.

**Figure 4.**
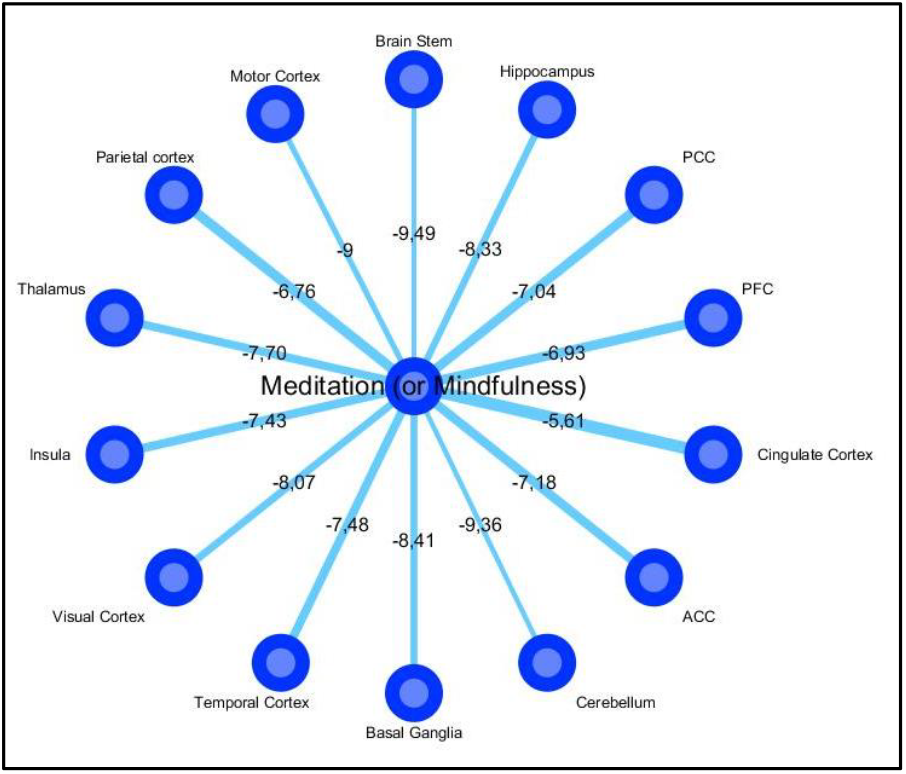
Mindfulness and brain area co-occurrences. The figure shows the most cited co-occurrences in the scientific literature of the terms “Meditation” or “Mindfulness” with the main brain areas. Significantly, the brain areas that are supposed to be involved in both meditation and consciousness had higher scores on the Jaccard Index, also indicated by thicker radii (the ln of the index is reported). [Data extracted with PubAtlas].

The involvement of four of these regions (i.e., the insula, ACC, PCC, and PFC), whose activity is considered to be extremely relevant in supporting both meditative and conscious states, is examined in the following paragraphs.

### 5.1. The role of the Insula and the ACC

There is evidence that during deep meditation the striatum, the left insula and the ACC are functionally active, whereas the lateral PFC and parietal cortex show reduced activation (Craigmyle, 2013; Hasenkamp et al., 2012; Hölzel et al., 2011; Posner et al., 2010; Tang et al., 2009; Tang and Posner, 2009). As we have seen, the ACC is thought to be a part of the “Internal awareness network” (Demertzi et al., 2013) and, together with the insula, is a crucial component of Seth’s interoceptive prediction model (Seth et al., 2012).

These two brain areas, which show structural alteration in long-term meditators (Craigmyle, 2013; Lazar et al., 2005), are also abundant in VENs (Cauda et al., 2014), whose deterioration has been associated with loss of emotional awareness and self-consciousness in patients with frontotemporal dementia (Seeley et al., 2007a; Seeley et al., 2006; Sturm et al., 2006). Within the predictive model framework, ACC activity seems to correlate with the probability of error prediction (Brown and Braver, 2005), as well as with the control of explorative behaviors (Aston-Jones and Cohen, 2005). Along with the MPFC, the ACC seems to play a prominent role in the evaluation of possible future scenarios (Ridderinkhof et al., 2004), which is in line with the “proactive brain” hypothesis. Moreover, the ACC is an essential part of the GW model (Dehaene et al., 1998).

### 5.2. The role of the PCC and PFC

The decrease in activity observed in the lateral PFC and parietal cortex during meditation on a focal object such as breath (Hölzel et al., 2011; Posner et al., 2010; Tang et al., 2009; Tang and Posner, 2009) is consistent with the hypothesis that these brain areas are involved in the “external awareness network” (Demertzi et al., 2013). On the basis of real-time neuro-feedback graph analyses, Garrison et al. (2013) showed that the mental states described by meditators as “undistracted awareness” or “effortless doing” correspond to PCC deactivation, whereas the mental states described as “distracted awareness” or “control” correspond to PCC activation. The PCC, which is part of Demertzi’s “internal awareness network”, is metabolically active in normal conscious states but often impaired in case of coma or vegetative state (Cauda et al., 2010; Cauda et al., 2009; Demertzi et al., 2013). It has therefore been proposed that PCC co-activation patterns could be reliable markers of consciousness modulation (Amico et al., 2014).

Thus, empirical data strongly suggest that the practice of meditation can cause both structural and functional alterations within the neural networks that promote and maintain consciousness. This phenomenon is more likely to occur for long-term meditators (Goleman, 1988; Shapiro, 2008) and might lead to a sort of “altered experience of time and space” (Berkovich-Ohana et al., 2013). This feeling might be associated with decreased activity in the PCC (Brewer et al., 2013). This view is probably also consistent with the mental state that long-term meditation practitioners, who have mastered meditative techniques, describe as “the mind observing itself” (i.e., the observation of thoughts in a detached and non-judgmental way). The Dalai Lama observes that something similar happens when we think of past experiences, even though in this case there is no temporal synchronicity between the one who thinks and what is thought of (Dalai Lama et al., 1991).

### 5.3 Unresolved issues and future directions

An important issue that still needs to be investigated is how long the practice of meditation has to continue before it starts to generate any appreciable neurophysiological alteration, and whether or not this change persists once the practice is interrupted. A related question is that of establishing a criterion in order to have a clear-cut distinction between subjects who can be assigned to the group of meditators and those who can be assigned to the group of non-meditators.

Thus far the scientific research has focused primarily on examining how meditation can affect the neurophysiology of long-term Buddhist practitioners, but future investigations are needed to study whether changes in neurophysiological parameters can also be observed in novices to this practice. Longitudinal studies should therefore be planned to measure the impact of meditation over time.

The scientific research should also address the topic of how meditation can influence the activation of the resting state network (Froeliger et al., 2012), as well as of other brain networks, such as the SN, the CEN, and the dorsal and ventral attentional systems. The relationship between the ability to control and maintain attention and the practice of meditation is of particular interest, given that long-term meditators seem to exhibit better skills in exploiting attentional resources than non-meditators. What is more, this ability may inhibit cognitive and emotional processes (i.e., rumination) that, in turn, could cause or exacerbate stress, anxiety, or depression (Brefczynski-Lewis et al., 2007). As a result, long-term meditators might show psycho-emotional stability and better attentional skills (Aftanas and Golosheykin, 2005). This mindset could bring about changes in their lifestyle, which could also positively affect their health and personality, as well as changes in the quality of conscious experience, in particular by enhancing the awareness of internal bodily states (Rubia, 2009). If that were the case, we would expect to observe alterations in both the dorsal and ventral attentional systems of those who practice meditation. Future investigations should therefore address this topic and find out whether both systems are affected in the same way or one is more affected than the other.

Research about this issue could produce interesting results. In fact, since consciousness and attention are closely intertwined, it is plausible that the impact of mindfulness meditation on consciousness might heavily depend on changes in the way of orienting and controlling attention. As we have seen, interoceptive attention is essentially involved in the mechanism that, according to the predictive models of the brain, underlies the experience of conscious presence (Seth et al., 2012). What is more, attentional processes play a fundamental role in the functional organization described in the GW theory.

Finally, there is the intriguing, albeit very speculative possibility that the brain areas involved in meditative practice could form a specific wide network in long-term meditators. There is indeed evidence to suggest that the practice of mindfulness meditation is associated with neuroplastic changes in the anterior cingulate cortex, insula, temporo-parietal junction, and fronto-limbic regions (Hölzel et al., 2011). These neuroplastic mechanisms might therefore strengthen some pathways and promote the generation of a self-sustaining process. This “mindfulness meditation network” might be composed of other smaller network structures (such as those related to the resting state, as well as the dorsal and ventral attentional systems) capable of generating a higher-order level of brain organization.

## 6. CONCLUSION

Mindfulness meditation is a method for training the mind that has been practiced in Eastern countries for more than two millennia, but has only quite recently attracted the attention of neuroscientists. In particular, the neuroscientific study of mindfulness meditation has aroused a great deal of interest in psychotherapeutic contexts and has inspired different cognitive approaches to stress reduction and mood disorders (Tang et al., 2015). There is in fact compelling evidence that meditative practices can significantly influence cognitive and emotional processes with various benefits on physical and mental health (Lutz et al., 2007; Soler et al., 2014; Tang et al., 2015).

A promising hypothesis suggested by this review is that some brain areas involved in meditation and consciousness could overlap, albeit partially. Such overlap encompasses the ACC, the insula, the PCC, some regions of the prefrontal cortex and the thalamus. As a result, the practice of meditation might somehow affect certain features of consciousness. In other words, the pattern of activation of brain areas that are thought to promote and maintain conscious states could exhibit typical differences. This would make the neuroscientific inquiry on meditation very worthwhile in order to better understand both the potential impact of meditative techniques on the brain and the neural counterpart of subjective experience.

What is more, this type of research is of crucial importance if mental training based on meditation has the potential to evolve into a standard procedure for therapeutic use (Tang et al., 2015). Therefore, the time is ripe for an integrative approach characterized by a wider theoretical framework in which meditation could be taken into account from the neurophysiological, psychological and behavioral perspectives.

## ACKNOWLEDGEMENTS

The authors would like to thank the Department of Psychology – University of Turin and the GCS-fMRI Research Group – Koelliker Hospital for their support and assistance with this research. Special thanks also go to Prof. Giuliano Geminiani and Dr Sergio Duca, whose advice and guidance are always invaluable.

## REFERENCES

Aftanas, L., Golosheykin, S., 2005. Impact of regular meditation practice on EEG activity at rest and during evoked negative emotions. The International journal of neuroscience 115, 893–909.

Allman, J.M., Tetreault, N.A., Hakeem, A.Y., Manaye, K.F., Semendeferi, K., Erwin, J.M., Park, S., Goubert, V., Hof, P.R., 2010. The von Economo neurons in frontoinsular and anterior cingulate cortex in great apes and humans. Brain structure & function 214, 495–517.

Allman, J.M., Tetreault, N.A., Hakeem, A.Y., Manaye, K.F., Semendeferi, K., Erwin, J.M., Park, S., Goubert, V., Hof, P.R., 2011. The von Economo neurons in the frontoinsular and anterior cingulate cortex. Annals of the New York Academy of Sciences 1225, 59–71.

Allman, J.M., Watson, K.K., Tetreault, N.A., Hakeem, A.Y., 2005. Intuition and autism: a possible role for Von Economo neurons. Trends in cognitive sciences 9, 367–373.

Amico, E., Gomez, F., Di Perri, C., Vanhaudenhuyse, A., Lesenfants, D., Boveroux, P., Bonhomme, V., Brichant, J.F., Marinazzo, D., Laureys, S., 2014. Posterior cingulate cortex-related co-activation patterns: a resting state FMRI study in propofol-induced loss of consciousness. PloS one 9, e100012.

Arzy, S., Thut, G., Mohr, C., Michel, C.M., Blanke, O., 2006. Neural basis of embodiment: distinct contributions of temporoparietal junction and extrastriate body area. The Journal of neuroscience: the official journal of the Society for Neuroscience 26, 8074–8081.

Aston-Jones, G., Cohen, J.D., 2005. An integrative theory of locus coeruleus-norepinephrine function: adaptive gain and optimal performance. Annual review of neuroscience 28, 403–450.

Awasthi, B., 2012. Issues and perspectives in meditation research: in search for a definition. Frontiers in psychology 3, 613.

Baars, B.J., 1988. A Cognitive Theory of Consciousness. Cambridge University Press.

Baars, B.J., Ramsoy, T.Z., Laureys, S., 2003. Brain, conscious experience and the observing self. Trends Neurosci 26, 671–675.

Bar, M., 2007. The proactive brain: using analogies and associations to generate predictions. Trends in cognitive sciences 11, 280–289.

Bar, M., Aminoff, E., Mason, M., Fenske, M., 2007. The units of thought. Hippocampus 17, 420–428.

Berkovich-Ohana, A., Dor-Ziderman, Y., Glicksohn, J., Goldstein, A., 2013. Alterations in the sense of time, space, and body in the mindfulness-trained brain: a neurophenomenologically-guided MEG study. Frontiers in psychology 4, 912.

Blanke, O., Landis, T., Spinelli, L., Seeck, M., 2004. Out-of-body experience and autoscopy of neurological origin. Brain: a journal of neurology 127, 243–258.

Blanke, O., Metzinger, T., 2009. Full-body illusions and minimal phenomenal selfhood. Trends in cognitive sciences 13, 7–13.

Blanke, O., Mohr, C., 2005. Out-of-body experience, heautoscopy, and autoscopic hallucination of neurological origin Implications for neurocognitive mechanisms of corporeal awareness and self-consciousness. Brain research. Brain research reviews 50, 184–199.

Blumenfeld, H., 2009. Epilepsy and consciousness, in: Laureys, S., Tononi, G. (Eds.), The Neurology of consciousness. Elsevier, Amsterdam, pp. 247–260.

Brefczynski-Lewis, J.A., Lutz, A., Schaefer, H.S., Levinson, D.B., Davidson, R.J., 2007. Neural correlates of attentional expertise in long-term meditation practitioners. Proceedings of the National Academy of Sciences of the United States of America 104, 11483–11488.

Bressler, S.L., Menon, V., 2010. Large-scale brain networks in cognition: emerging methods and principles. Trends in cognitive sciences 14, 277–290.

Brewer, J.A., Garrison, K.A., Whitfield-Gabrieli, S., 2013. What about the “Self” is Processed in the Posterior Cingulate Cortex? Frontiers in human neuroscience 7, 647.

Brown, J.W., Braver, T.S., 2005. Learned predictions of error likelihood in the anterior cingulate cortex. Science (New York, N.Y.) 307, 1118–1121.

Brugger, P., 2006. From phantom limb to phantom body: Varieties of extracorporeal awareness, in: Knoblich, G., Thornton, I.M., Grosjean, M., Shiffrar, M. (Eds.), Human Body Perception From the Inside Out. Oxford University Press, pp. 171–209.

Butti, C., Sherwood, C.C., Hakeem, A.Y., Allman, J.M., Hof, P.R., 2009. Total number and volume of Von Economo neurons in the cerebral cortex of cetaceans. The Journal of comparative neurology 515, 243–259.

Cauda, F., D'Agata, F., Sacco, K., Duca, S., Geminiani, G., Vercelli, A., 2011. Functional connectivity of the insula in the resting brain. NeuroImage 55, 8–23.

Cauda, F., Geminiani, G., D'Agata, F., Sacco, K., Duca, S., Bagshaw, A.P., Cavanna, A.E., 2010. Functional Connectivity of the Posteromedial Cortex. PloS one 5, e13107.

Cauda, F., Geminiani, G.C., Vercelli, A., 2014. Evolutionary appearance of von Economo's neurons in the mammalian cerebral cortex. Frontiers in human neuroscience 8, 104.

Cauda, F., Micon, B.M., Sacco, K., Duca, S., D'Agata, F., Geminiani, G., Canavero, S., 2009. Disrupted intrinsic functional connectivity in the vegetative state. Journal of Neurology, Neurosurgery & Psychiatry 80, 429–431.

Cauda, F., Torta, D.E., Sacco, K., D'Agata, F., Geda, E., Duca, S., Geminiani, G., Vercelli, A., 2013. Functional anatomy of cortical areas characterized by Von Economo neurons. Brain Structure and Function 218, 1–20.

Cauda, F., Torta, D.M.E., Sacco, K., Geda, E., D'Agata, F., Costa, T., Duca, S., Geminiani, G., Amanzio, M., 2012. Shared “Core” Areas between the Pain and Other Task-Related Networks. PloS one 7, e41929.

Cavanna, A.E., Shah, S., Eddy, C.M., Williams, A., Rickards, H., 2011. Consciousness: a neurological perspective. Behavioural neurology 24, 107–116.

Chiesa, A., Serretti, A., 2010. A systematic review of neurobiological and clinical features of mindfulness meditations. Psychological Medicine 40, 1239–1252.

Chiesa, A., Serretti, A., 2011. Mindfulness based cognitive therapy for psychiatric disorders: a systematic review and meta-analysis. Psychiatry research 187, 441–453.

Christoff, K., Gordon, A.M., Smallwood, J., Smith, R., Schooler, J.W., 2009. Experience sampling during fMRI reveals default network and executive system contributions to mind wandering. Proceedings of the National Academy of Sciences 106, 8719–8724.

Craig, A.D., 2002. How do you feel? Interoception: the sense of the physiological condition of the body. Nature reviews. Neuroscience 3, 655–666.

Craig, A.D., 2004. Human feelings: why are some more aware than others? Trends in cognitive sciences 8, 239–241.

Craig, A.D., 2009a. Emotional moments across time: a possible neural basis for time perception in the anterior insula. Philosophical transactions of the Royal Society of London. Series B, Biological sciences 364, 1933–1942.

Craig, A.D., 2009b. How do you feel - now? The anterior insula and human awareness. Nature reviews. Neuroscience 10, 59–70.

Craigmyle, N.A., 2013. The beneficial effects of meditation: contribution of the anterior cingulate and locus coeruleus. Frontiers in psychology 4, 731.

Critchley, H., Seth, A., 2012. Will studies of macaque insula reveal the neural mechanisms of self-awareness? Neuron 74, 423–426.

Critchley, H.D., Wiens, S., Rotshtein, P., Ohman, A., Dolan, R.J., 2004. Neural systems supporting interoceptive awareness. Nat Neurosci 7, 189–195.

Dalai Lama, Thurman, R.A.F., Gardner, H.E., Goleman, D., 1991. MindScience: An East-West Dialogue. Wisdom Publications.

Dayan, P., Hinton, G.E., Neal, R.M., Zemel, R.S., 1995. The Helmholtz machine. Neural computation 7, 889–904.

Dehaene, S., Changeux, J.P., 2011. Experimental and theoretical approaches to conscious processing. Neuron 70, 200–227.

Dehaene, S., Kerszberg, M., Changeux, J.P., 1998. A neuronal model of a global workspace in effortful cognitive tasks. Proc. Natl. Acad. Sci. USA 95.

Dehaene, S., Naccache, L., 2001. Towards a cognitive neuroscience of consciousness: basic evidence and a workspace framework. Cognition 79, 1–37.

Demertzi, A., Soddu, A., Laureys, S., 2013. Consciousness supporting networks. Current opinion in neurobiology 23, 239–244.

Devinsky, O., Feldmann, E., Burrowes, K., Bromfield, E., 1989. AUtoscopic phenomena with seizures. Archives of Neurology 46, 1080–1088.

Farb, N.A., Segal, Z.V., Anderson, A.K., 2013. Mindfulness meditation training alters cortical representations of interoceptive attention. Social cognitive and affective neuroscience 8, 15–26.

Farb, N.A.S., Anderson, A.K., Mayberg, H., Bean, J., McKeon, D., Segal, Z.V., 2010. Minding one's emotions: mindfulness training alters the neural expression of sadness. Emotion 10, 25–33.

Farb, N.A.S., Segal, Z.V., Mayberg, H., Bean, J., McKeon, D., Fatima, Z., Anderson, A.K., 2007. Attending to the present: mindfulness meditation reveals distinct neural modes of self-reference. Social cognitive and affective neuroscience 2, 313–322.

Flynn, F.G., 1999. Anatomy of the insula functional and clinical correlates. Aphasiology 13, 55–78.

Friston, K., 2010. The free-energy principle: a unified brain theory? Nature reviews. Neuroscience 11, 127–138.

Friston, K., 2012. The history of the future of the Bayesian brain. NeuroImage 62, 1230–1233.

Friston, K., Kilner, J., Harrison, L., 2006. A free energy principle for the brain. Journal of physiology, Paris 100, 70–87.

Froeliger, B., Garland, E.L., Kozink, R.V., Modlin, L.A., Chen, N.K., McClernon, F.J., Greeson, J.M., Sobin, P., 2012. Meditation-State Functional Connectivity (msFC): Strengthening of the Dorsal Attention Network and Beyond. Evidence-based complementary and alternative medicine: eCAM 2012, 680407.

Garrison, K.A., Santoyo, J.F., Davis, J.H., Thornhill, T.A.t., Kerr, C.E., Brewer, J.A., 2013. Effortless awareness: using real time neurofeedback to investigate correlates of posterior cingulate cortex activity in meditators' self-report. Frontiers in human neuroscience 7, 440.

Goldin, P., Ziv, M., Jazaieri, H., Hahn, K., Gross, J.J., 2013. MBSR vs aerobic exercise in social anxiety: fMRI of emotion regulation of negative self-beliefs. Social cognitive and affective neuroscience 8, 65–72.

Goleman, D., 1988. The Meditative Mind: The Varieties of Meditative Experience. Tarcher.

Grafton, S.T., Hazeltine, E., Ivry, R., 1995. Functional anatomy of motor sequence learning in humans. J. Cognit. Neurosci. 7, 497–510.

Gregory, R.L., 1980. Perceptions as Hypotheses. Philosophical Transactions of the Royal Society of London. B, Biological Sciences 290, 181–197.

Hasenkamp, W., Wilson-Mendenhall, C.D., Duncan, E., Barsalou, L.W., 2012. Mind wandering and attention during focused meditation: A fine-grained temporal analysis of fluctuating cognitive states. NeuroImage 59, 750–760.

Hölzel, B.K., Lazar, S.W., Gard, T., Schuman-Olivier, Z., Vago, D.R., Ott, U., 2011. How Does Mindfulness Meditation Work? Proposing Mechanisms of Action From a Conceptual and Neural Perspective. Perspect. Psychol. Sci. 6, 22.

Kabat-Zinn, J., 2003. Mindfulness-Based Interventions in Context: Past, Present, and Future. Clinical Psychology: Science and Practice 10, 144–156.

Kersten, D., Mamassian, P., Yuille, A., 2004. Object Perception as Bayesian Inference. Annual Review of Psychology 55, 271–304.

Knill, D.C., Pouget, A., 2004. The Bayesian brain: the role of uncertainty in neural coding and computation. Trends Neurosci 27, 712–719.

Kozasa, E.H., Sato, J.R., Lacerda, S.S., Barreiros, M.A., Radvany, J., Russell, T.A., Sanches, L.G., Mello, L.E., Amaro, E., Jr., 2012. Meditation training increases brain efficiency in an attention task. NeuroImage 59, 745–749.

Laureys, S., 2005. Death, unconsciousness and the brain. Nature reviews. Neuroscience 6, 899–909.

Laureys, S., Boly, M., 2008. The changing spectrum of coma. Nature clinical practice. Neurology 4, 544–546.

Laureys, S., Owen, A.M., Schiff, N.D., 2004. Brain function in coma, vegetative state, and related disorders. The Lancet. Neurology 3, 537–546.

Lazar, S.W., Kerr, C.E., Wasserman, R.H., Gray, J.R., Greve, D.N., Treadway, M.T., McGarvey, M., Quinn, B.T., Dusek, J.A., Benson, H., Rauch, S.L., Moore, C.I., Fischl, B., 2005. Meditation experience is associated with increased cortical thickness. Neuroreport 16, 1893–1897.

Lee, T.S., Mumford, D., 2003. Hierarchical Bayesian inference in the visual cortex. Journal of the Optical Society of America. A, Optics, image science, and vision 20, 1434–1448.

Lutz, A., Dunne, J.D., Davidson, R.J., 2007. Meditation and the neuroscience of consciousness, in: Zelazo, P.D., Moscovitch, M., Thompson, E. (Eds.), Cambridge Handbook of Consciousness. Cambridge, pp. 19–497.

Medford, N., Critchley, H.D., 2010. Conjoint activity of anterior insular and anterior cingulate cortex: awareness and response. Brain structure & function 214, 535–549.

Menon, V., Uddin, L., 2010. Saliency, switching, attention and control: a network model of insula function. Brain Structure and Function 214, 655–667.

Merkes, M., 2010. Mindfulness-based stress reduction for people with chronic diseases. Australian Journal of Primary Health 16, 200–210.

Mesulam, M.M., 1998. From sensation to cognition. Brain: a journal of neurology 121, 1013–1052.

Metzinger, T., Gallese, V., 2003. The emergence of a shared action ontology: building blocks for a theory. Consciousness and cognition 12, 549–571.

Mullette-Gillman, O.D.A., Huettel, S.A., 2009. Neural substrates of contingency learning and executive control: dissociating physical, valuative, and behavioral changes. Frontiers in human neuroscience 3, 23.

Nani, A., Seri, A., Cavanna, A.E., 2013. Consciousness and Neuroscience, in: Cavanna, A.E., Nani, A., Blumenfeld, H., Laureys, S. (Eds.), Neuroimaging of Consciousness. Springer Verlag, Berlin, pp. 3–21.

Pacherie, E., 2008. The phenomenology of action: A conceptual framework. Cognition 107, 179–217.

Palaniyappan, L., Liddle, P.F., 2012. Does the salience network play a cardinal role in psychosis? An emerging hypothesis of insular dysfunction. Journal of psychiatry & neuroscience: JPN 37, 17–27.

Plum, F., Posner, J.B., 1980. The diagnosis of stupor and coma, 3rd ed. Davis, Philadelphia.

Posner, M.I., Rothbart, M.K., Rueda, M.R., Tang, Y., 2010. Training effortless attention, in: Bruya, B. (Ed.), Effortless Attention: A New Perspective in the Cognitive Science of Attention and Action. Mit Press.

Ridderinkhof, K.R., van den Wildenberg, W.P., Segalowitz, S.J., Carter, C.S., 2004. Neurocognitive mechanisms of cognitive control: the role of prefrontal cortex in action selection, response inhibition, performance monitoring, and reward-based learning. Brain and cognition 56, 129–140.

Roessler, J., Eilan, N., 2003. Agency and Self-Awareness: Issues in Philosophy and Psychology. Oxford University Press.

Rubia, K., 2009. The neurobiology of Meditation and its clinical effectiveness in psychiatric disorders. Biological psychology 82, 1–11.

Sahraie, A., Weiskrantz, L., Barbur, J.L., Simmons, A., Williams, S.C., Brammer, M.J., 1997. Pattern of neuronal activity associated with conscious and unconscious processing of visual signals. Proceedings of the National Academy of Sciences of the United States of America 94, 9406–9411.

Samuel, G., 2014. The contemporary mindfulness movement and the question of nonself1. Transcultural psychiatry.

Sato, J.R., Kozasa, E.H., Russell, T.A., Radvany, J., Mello, L.E., Lacerda, S.S., Amaro, E., Jr., 2012. Brain imaging analysis can identify participants under regular mental training. PloS one 7, e39832.

Seeley, W.W., 2008. Selective functional, regional, and neuronal vulnerability in frontotemporal dementia. Current opinion in neurology 21, 701–707.

Seeley, W.W., Allman, J.M., Carlin, D.A., Crawford, R.K., Macedo, M.N., Greicius, M.D., Dearmond, S.J., Miller, B.L., 2007a. Divergent social functioning in behavioral variant frontotemporal dementia and Alzheimer disease: reciprocal networks and neuronal evolution. Alzheimer disease and associated disorders 21, S50–57.

Seeley, W.W., Carlin, D.A., Allman, J.M., Macedo, M.N., Bush, C., Miller, B.L., DeArmond, S.J., 2006. Early frontotemporal dementia targets neurons unique to apes and humans. Annals of Neurology 60, 660–667.

Seeley, W.W., Menon, V., Schatzberg, A.F., Keller, J., Glover, G.H., Kenna, H., Reiss, A.L., Greicius, M.D., 2007b. Dissociable intrinsic connectivity networks for salience processing and executive control. The Journal of neuroscience: the official journal of the Society for Neuroscience 27, 2349–2356.

Seth, A.K., Suzuki, K., Critchley, H.D., 2012. An interoceptive predictive coding model of conscious presence. Frontiers in psychology 2.

Shallice, T., 1988. From Neuropsychology to Mental Structure. Cambridge University Press.

Shapiro, D.H., 2008. Meditation: Self-Regulation Strategy and Altered State of Consciousness. Aldine De Gruyter, NY.

Siegel, R.D., Germer, C.K., Olendzki, A., 2008. Mindfulness: What Is It? Where Did It Come From?, in: Didonna, F. (Ed.), Clinical Handbook of Mindfulness. Springer, New York.

Soler, J., Cebolla, A., Feliu-Soler, A., Demarzo, M.M.P., Pascual, J.C., Baños, R., García-Campayo, J., 2014. Relationship between Meditative Practice and Self-Reported Mindfulness: The MINDSENS Composite Index. PloS one 9, e86622.

Steriade, M., 1996a. Arousal: revisiting the reticular activating system. Science (New York, N.Y.) 272, 225–226.

Steriade, M., 1996b. Awakening the brain. Nature 383, 24–25.

Stimpson, C.D., Tetreault, N.A., Allman, J.M., Jacobs, B., Butti, C., Hof, P.R., Sherwood, C.C., 2011. Biochemical specificity of von Economo neurons in hominoids. American journal of human biology: the official journal of the Human Biology Council 23, 22–28.

Sturm, V.E., Rosen, H.J., Allison, S., Miller, B.L., Levenson, R.W., 2006. Self-conscious emotion deficits in frontotemporal lobar degeneration. Brain: a journal of neurology 129, 2508–2516.

Tang, Y.Y., Holzel, B.K., Posner, M.I., 2015. The neuroscience of mindfulness meditation. Nature reviews. Neuroscience 16, 213–225.

Tang, Y.Y., Ma, Y., Fan, Y., Feng, H., Wang, J., Feng, S., Lu, Q., Hu, B., Lin, Y., Li, J., Zhang, Y., Wang, Y., Zhou, L., Fan, M., 2009. Central and autonomic nervous system interaction is altered by shortterm meditation. Proceedings of the National Academy of Sciences of the United States of America 106, 8865–8870.

Tang, Y.Y., Posner, M.I., 2009. Attention training and attention state training. Trends in cognitive sciences 13, 222–227.

Tang, Y.Y., Rothbart, M.K., Posner, M.I., 2012. Neural correlates of establishing, maintaining, and switching brain states. Trends in cognitive sciences 16, 330–337.

Taylor, K.S., Seminowicz, D.A., Davis, K.D., 2009. Two systems of resting state connectivity between the insula and cingulate cortex. Human brain mapping 30, 2731–2745.

Thera, N., 1962. Heart of Buddhist Meditation Buddhist Publication Society, Kandy, Sri Lanka.

Torta, D.M., Cauda, F., 2011. Different functions in the cingulate cortex, a meta-analytic connectivity modeling study. NeuroImage 56, 2157–2172.

van den Heuvel, M.P., Mandl, R.C., Kahn, R.S., Hulshoff Pol, H.E., 2009. Functionally linked resting-state networks reflect the underlying structural connectivity architecture of the human brain. Human brain mapping 30, 3127–3141.

Van Oudenhove, L., Vandenberghe, J., Dupont, P., Geeraerts, B., Vos, R., Bormans, G., Van Laere, K., Fischler, B., Demyttenaere, K., Janssens, J., Tack, J., 2009. Cortical deactivations during gastric fundus distension in health: visceral pain-specific response or attenuation of 'default mode' brain function? A H215O-PET study. Neurogastroenterology & Motility 21, 259–271.

von Economo, C., 1926. Eine neue Art Spezialzellen des Lobus cinguli und Lobus insulae. Z. GESAMTE NEUROL. PSYCHIATR 100, 706–712.

von Economo, C., 1927. L'architecture cellulaire normale de l'ecorce cérébrale. Paris: Masson.

von Economo, C., Koskinas, G.N., 1925. Die cytoarchitektonik der hirnrinde des erwachsenen menschen. Berlin: Verlag von Julius Springer.

Weiskrantz, L., 1997. Consciousness Lost and Found: A Neuropsychological Exploration. Oxford University Press, New York.

Zeman, A., 2001. Consciousness. Brain: a journal of neurology 124, 1263–1289.

